# The interplay of recombination landscape, a transposable element and population history in European populations of *Chironomus riparius*

**DOI:** 10.1101/2024.04.04.588075

**Authors:** Laura C. Pettrich, Robert King, Linda M. Field, Ann-Marie Waldvogel

## Abstract

Genome resolution is often constrained for non-model species. This can be challenging for population genomic studies as estimations are highly dependent on the quality of the reference genome. This is the case for population history inferences where accuracy relies on the correct detection of single-nucleotide polymorphisms (SNPs) and accurate recombination rates. Here, we utilize a novel long-read genome assembly of *Chironomus riparius* with high resolution at a chromosome-scale and reanalyse Illumina resequencing data of five European populations. With the model MSMC2, new population demographies were inferred and compared to an older study based on a fragmented genome. Assembly contiguity and completeness led to an increase in accuracy of past demography, suggesting the onset of divergence of an ancestral population between late Pleistocene and early Holocene. These estimates are additionally supported by paleotemperature data, which reveal significant climate shifts in Central Europe during these times. Recombination severely influences population history estimates. With the new reference genome, it was possible to resolve the recombination landscape on the genome-wide scale across different populations using the tool iSMC. Recombination and the dispersal of transposable elements (TEs) in the genome are suspected to influence each other. One TE, known as *Cla*-element, is suspected to be involved in the divergence of the different *C. riparius* populations. As it seems to be highly dynamic in the genome, its potential impact on the recombination landscape was explored. No global pattern could be detected which demonstrated a higher resistance of the recombination landscape to the impact of repetitive elements on genome integrity.

## Significance

Population history models can be influenced by recombination variation. Based on a new genome assembly the recombination variation of different European *Chironomus riparius* populations was explored and compared to a transposable element suspected of driving population divergence. Resolving influencers of recombination variation have the potential to improve population demography studies.

## Introduction

Patterns of population growth or decline (Li & Durbin 2011; Schiffels & Durbin 2014) and admixture among populations (Luikart et al. 2003) can be read out from genomic data. Studying evolution and demography of natural populations is facilitated by whole-genome resequencing data of multiple individuals (Waldvogel et al. 2020; Bourgeois & Warren 2021).

Thus, a rise in population genomic studies can be associated with the increasing availability of high-quality reference genomes due to advances in long-read sequencing technologies (Guiglielmoni et al. 2023; The Darwin Tree of Life Project Consortium 2022; Blaxter et al. 2022). *Chironomus riparius*, a non-biting midge, commonly known as the harlequin fly and distributed in Europe, has been the focus of many molecular genomic studies (e.g. Oppold et al. 2017 and Waldvogel et al. 2018), with work dating back to 1984 (Hägele 1984). Polytene chromosomes in their salivary glands became the basis to study the species’ genome structure and trace banding patterns of heterochromatic regions in the genome during crossings with related *Chironomus* species (Schmidt 1981; Hankeln & Schmidt 1987; Bovero et al. 2002; Hägele 1984). These early studies started to document the species’ genome and many molecular genetic works followed, applying different methodology and sequencing strategies (Schreiber & Pfenninger 2021; Oppold et al. 2017; Schmidt et al. 2020). In this respect, the *Chironomus riparius* genome project also reflects the continuous advancement of sequencing technology and assembly strategy. With this study, we present a high-quality genome assembly for the species that reaches chromosome scale, marking a new milestone for future studies with this model system. Furthermore, this genomic resource now allows for a more accurate estimation of population history and reconstruction of the recombination landscape.

For the inference of population demography, sequential Markovian coalescent models (SMC) (Li & Durbin 2011; Schiffels & Wang 2020; Wilton et al. 2015) trace back mutation and recombination events. In this, a hidden Markov model (HMM) (Wiuf & Hein 1999) in a sequential Markovian coalescence (SMC) framework (Marjoram & Wall 2006; McVean & Cardin 2005) derives hidden states together with its transition rates and estimates coalescence times along the sequence. Applying genomic approaches like these to a broader taxonomic spectrum can help to revisit traditional and long-standing questions about the evolution of genetic diversity and genomic mechanisms of biodiversification (Blaxter et al. 2022).The Lewontin’s paradox (Lewontin 1974), one such long-standing open question of biology, suggests that neutral genetic diversity remains stable between species despite varying population census sizes with many orders of magnitude differences (Charlesworth & Jensen 2022; Roberts 2015). It has been discussed whether natural selection causes this kind of equilibrium between population size and heterozygosity. Advantageous variations that are passed down to the next generation, together with hitchhiking genes, might cause a reduction in diversity of certain genomic regions (Roberts 2015). Eradicating disadvantageous variations may have a similar effect (Roberts 2015). Determining the molecular cause of Lewontin’s paradox remains challenging due to the convolution of genomic signatures that result from selection and/or demographic changes (Charlesworth & Jensen 2022). It is thus crucial to explore aspects that can help to improve approaches that allow for a more accurate inference of demographic histories and resolve periods of expansion in population size (Charlesworth & Jensen 2022), in particular for a broader range of taxa that also differ in their life history traits (Sellinger et al. 2020).

Gaining a comprehensive understanding of the crucial parameters, such as recombination and mutation rates, is essential to accurately infer population histories. Mutation rates have been shown to depend on certain environmental conditions (Waldvogel & Pfenninger 2021) and recombination rates show variations on a chromosomal scale differing tremendously between species (Nachman 2002). Genomic regions with low recombination often show lower nucleotide diversity (Nachman 2002). When inferring population history, it may be necessary to exclude heterogeneous regions with five-fold differences in recombination rates, as false estimates in past effective population sizes have been documented (Sellinger et al. 2021). On top of this, the landscape of a genome is severely impacted by genetic markers, with transposable elements (TEs) playing a key role in genomic rearrangements (Montgomery et al. 1991). Despite this knowledge on the genomic impact of TEs, how they influence the genome-wide landscape of recombination is still poorly understood (Kent et al. 2017). It appears that TEs are often associated with low recombing regions and the extent of interaction hints at a possible contribution of TE insertions to recombination rate variation (Kent et al. 2017). Repetitive regions are generally expected to have lower recombination rates (Kent et al. 2017), especially in centromere regions and regions with large regions of heterochromatin, where recombination is known to be reduced, while it can be elevated in telomere regions (Kent et al. 2017; Nachman 2002). Recombination hotspots are known to occur in GC-rich regions (Marsolier-Kergoat & Yeramian 2009). A high-quality genome gives access to regions that are known to vary in recombination rates (Pollard et al. 2018; Nachman 2002) which is expected to make demographic estimates more robust, as subsequently more information uncovering genealogy becomes available. Recombination is an important factor to consider when using SMC models, as they utilise ancestral recombination events (i.e ancestral recombination graphs (ARG)) to infer population history (McVean & Cardin 2005; Marjoram & Wall 2006; Mather et al. 2020). As such, it becomes important to look at the whole genomic landscape to investigate population history (Soni et al. 2024). Many population demography models were developed using genome data of humans or model organisms like *Drosophila melanogaster* and *Arabidopsis thaliana* (Li & Durbin 2011; Schiffels & Durbin 2014) with the result that these models were built to rely on high-quality reference genomes with known genomic characteristics like ploidy, maps of recombination and a mutational landscape. Studying population genomics in a broader taxonomic spectrum, beyond genomically well-investigated model species, however, often involves reference genomes of lower quality and more complex genomic characteristics, such as polyploidy or life cycles with reduced sexual reproduction what affects the frequency of meiosis, i.e. recombination. To account for these challenges, model refinements are in development to better account for violations of SMC models, as for example dormancy (Sellinger et al. 2020). The importance of considering markers like TEs, microsatellites and DNA methylation in demographic inferences are also being explored in model organisms and yield improved results in assessing structuring events like bottlenecks or expansions (Sellinger et al. 2023). Genomic markers with elevated mutation rates have been shown to improve temporal resolution of recent evolutionary estimates (Sellinger et al. 2023) hence showing the importance of including not only Single-Nucleotide Polymorphisms (SNP).

The genome of *Chironomus riparius* contains a ‘special’ transposable element, known as the *Cla*-element, that has attracted attention in molecular genetic as well as population genomic studies and is hypothesized to have been involved in the speciation process of the species (Schmidt 1984; Oppold et al. 2017). The *Cla*-element has a consensus sequence of 120 bp (Schmidt 1981) and can be found in repetitive clusters (Hankeln et al. 2011) or as monomeric repeats (Hankeln & Schmidt 1987). The element has been shown to drive genomic patterns of population divergence in *C. riparius* (Oppold et al. 2017). Genome-wide patterns of Cla-element insertions were found to vary among natural populations, which, together with the finding of *Cla* transcripts and heterozygous monomer insertions, led to the assumption of ongoing transposition activity of the element (Oppold et al. 2017). In the model organism *Drosophila*, piRNA and siRNA pathways are involved in the control of different TEs (Mérel et al. 2020) and it seems that piRNA-mediated silencing could be ancestral in arthropods (Lewis et al. 2018). In the grasshopper *Angaracris rhodopa* lower piRNA concentrations are suspected to be involved in the expansion of its genome due to higher TE activity (Liu et al. 2022). In the Antarctic midge chironomid *Belgica antarctica* piRNAs have been documented which might explain low TE abundance (Kozeretska et al. 2022; Kelley et al. 2014). As such, regulation of TEs may also play an important role in the genome evolution of *C. riparius*. Compared to its sister species *C. piger*, *C. riparius* has a larger genome (Hankeln & Schmidt 1987; Schaefer & Schmidt 1981). A lack of *Cla*-element suppression could be the explanation for the spread of this element into chromosomal arms of *C. riparius* compared to its sister species *C. piger* Crossing of different populations resulted in less fit offspring and it was observed that heterozygous *Cla*-element patterns lead to incorrect synapsis of homologous chromosomes (Oppold et al. 2017). The *Cla*-element seems to cause Dobzhansky-Muller incompatibilities (DMI) between the investigated populations, showing the evolution of an endogenous genetic barrier (Oppold et al. 2017). This TE is known to form hairpin loop formations (Schmidt 1984; Israelewski 1983) which might cause ectopic recombination events (Oppold et al. 2017). These different aspects indicate a particular speciation potential of the *Cla*-element in *C. riparius*, making it particularly relevant to investigate its impact on the recombination landscape of the diverging populations.

This study aims to investigate the interplay of recombination, the *Cla*-element as a candidate TE with high potential to impact in recombination processes and population history of the non-biting midge *C. riparius*. Utilising a new long-read reference genome, this is the first attempt to understand variation of the recombination landscape in different populations of the harlequin fly while comparing them to the pattern of a particular transposable element (TE)-the *Cla*-element. For this species and the exact same populations, a previous study has been investigating the population history based on a fragmented genome draft (Waldvogel et al. 2018). Direct comparison of the inferred demographic histories to Waldvogel et al. (2018) provides direct insight into the importance and impact of the resolution of a reference genome on SMC studies. To estimate the population demography, we used the Multiple Markovian Coalescent (MSMC2) model, which uses a hidden Markov model to estimate genealogies along the sequences (Schiffels & Wang 2020). Further correlating the demographic estimates with climate history models (Karger et al. 2021), we outline how such genomic data can help to understand biogeographic histories and how additional data sources, here climate data, can serve as validation stringency when interpreting genomically inferred demographic histories. We also test the hypothesis that recombination landscape and distribution of TEs (here the *Cla*-element) are correlated and population-specific.

## Results

### Influence of improved genome assembly on inferred population history

We mapped the Illumina reads of five European populations of *C. riparius*, from Hesse in Germany (MG), Rhône-Alpes (MF) and Lorraine (NMF) in France, Piemont in Italy (SI) and Andalusia in Spain (SS), to the novel high-quality reference genome assembly (mapping statistics can be found in Supplementary Table S3). The assembly resolves all four chromosomes of *C. riparius* with ten remaining unplaced scaffolds. Genomes of two endosymbionts, *Enterobacter* sp. and *Wolbachia* sp. were assembled from the metafraction of the data. The *C. riparius*’ assembly spans 192 Mb with N50 of 59 Mb (Table 1, Fig. 1A). The assembly’s completeness when compared to the single ortholog database of Diptera (diptera_odb10, n=3285) scores 92.6 % complete, 1.8 % duplicated, 1.3 % fragmented and 6.1 % missing BUSCO genes.

**Figure 1:**
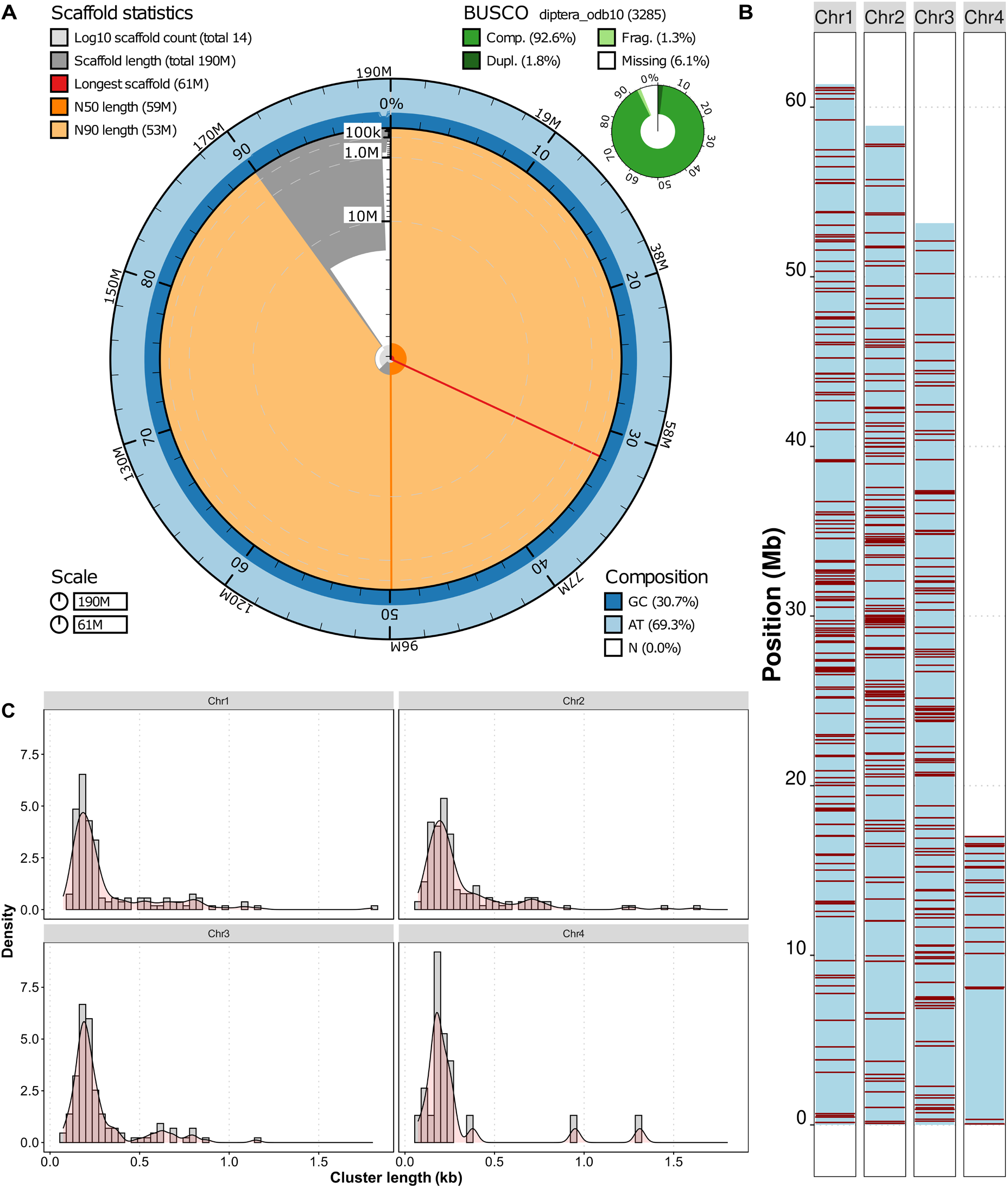
A) Snail plot summarising the assembly statistics. Scaffold statistics can be found on the top right. The longest scaffold (i.e. chromosome 1) is marked in red. The N50 value is marked in orange and the N90 value in pale orange. The GC and AT composition are in dark and light blue. A BUSCO analysis was performed against the diptera_odb10 database and values on completeness, fragmentation, duplication, and missing genes can be found on the top right. B) Plot of the four chromosomes with the total number of *Cla*-elements found by MELT across all populations. Red lines indicate *Cla*-elements. C) Density plots showing the distribution of the different lengths of *Cla*-element clusters (in kb) for each chromosome.

**Table 1:** Assembly statistics of the genome.

|  |  |
| --- | --- |
| Assembly size | 191,837,449 bp |
| N50 | 58,906,861 bp |
| GC content | 30.7 % |
| No. of repeats | 40473 |
| No. of protein-coding genes | 15439 |
| BUSCO (diptera_odb10) | C: 92.6 % [S: 90.8 %, D: 1.8 %], F: 1.3 %, M: 6.1 %, n: 3285 |
| Longest chromosome | 61,357,614 bp |
| No. of chromosomes | 4 |
| No. of unplaced scaffolds | 10 |

Chromosome numbers were ranked with descending length, with chromosome 1 being the longest scaffold of 61 Mb. The genome-wide GC-content was estimated to be 30.7 %. The assembly had a repeat content of 15.85 %. In total 15439 protein-coding genes were annotated. *Cla*-elements were detected using RepeatMasker and MELT and a total number of 433 *Cla*-elements insertions were detected (1492 monomeric repeats were detected with RepeatMasker) across the four chromosomes integrating all populations (Figure 1B), with 8 insertions on the unplaced scaffolds. Cla-elements are distributed between chromosomes with 148 on chromosome 1, 143 on chromosome 2, 119 on chromosome 3 and 20 on chromosome 4. The distribution of the different *Cla*-element cluster lengths (in kb) for each chromosome can be found in Figure 1C.

The Multiple Sequentially Markovian Coalescent (MSMC2) model was used to infer the population history of the five European *C. riparius* populations based on the novel high-quality genome assembly (details on the input files in Supplementary Table S4). We assessed admixture between populations (Figure 2A) and the history of effective population size (Figure 2B). The first five and the last time intervals of the estimation were excluded from the analysis to avoid uncertainties caused by the model due to overfitting. To improve the estimation robustness of the recent time horizon, we determined the time to the most recent common ancestor (tMRCA) of the individuals of a population by integrating a mean heterozygosity of 0.0083 and an informative mean haplotype length (MHL) of 6,023 bases. The tMRCA was determined to be 6,092 generations, when using the mean recombination rate of *C. riparius* (1.36 cM/Mb) or, alternatively, 3,953 generations when applying the recombination rate of *D. melanogaster* (2.1 cM/Mb) as applied in Waldvogel et al. (2018). Considering the species-specific recombination rate, informative time intervals range from 6,100 to 7,400,000 generations. When translating these coalescence times in years, the tMRCA in generations was multiplied with the mean generation time of the respective population (referring to estimates reported in Oppold et al. 2016) which resulted in a tMRCA of 609 years (see also Supplementary Table S6). After incorporating the generation time of each population into the inferred population history, the coalescence estimates of the ancient past converged, supported by a high relative cross-coalescence rate (rCCR) (Figure 2A) indicative for good admixture between populations. This admixture persisted until a peak 20,000 years ago (Figure 2B). Subsequent to the peak, in the more recent past, the effective population size experienced a decline, leading to a dispersal of the population-specific estimates. Concurrently, there was a reduction in rCCR estimated. The rCCR ranges between 0 and 1 and every value above one is considered an artefact caused by the model. A high rCCR indicates a high admixture between populations which can be found for our populations earlier than 10,000 years ago. Over time the admixture decreased and when the rCCR was below 0.5, the populations are estimated to have split into subpopulations. This process was estimated to have started approximately 10,000 years ago. We compared the inferred population demography to paleoclimate models from the CHELSA database, to assess whether certain temperature events could be responsible for the formation of different subpopulations (Figure 2C-F). By reviewing the annual mean temperature of the population’s habitat over time (Figure 2C) a gradient can be found. Hesse and Lorraine are the coldest regions, followed by Rhône-Alpes and Piemont and the warmest region has been Andalusia. Shifting the focus from only the population’s habitat to the European region (Figure 2D-F) a warming of all of Europe can be observed. The Last-Glacial Maximum (LGM) was 22,000 years before present (22k-BP) where many regions had a mean temperature lower than 0 °C (Figure 2F). In the latest period, 1k-BP (Figure 2D), most of the regions were warmer with only mountain ranges and the far North showing mean temperatures lower than 0 °C. The historical temperature estimates north of the alps indicate a shift from relatively cold temperatures (-15 °C to 5 °C) to moderate temperatures (0 °C to 15 °C). According to the model, southern Europe exhibited a consistently moderate climate (5 °C to 20 °C) starting from the earliest time point.

**Figure 2:**
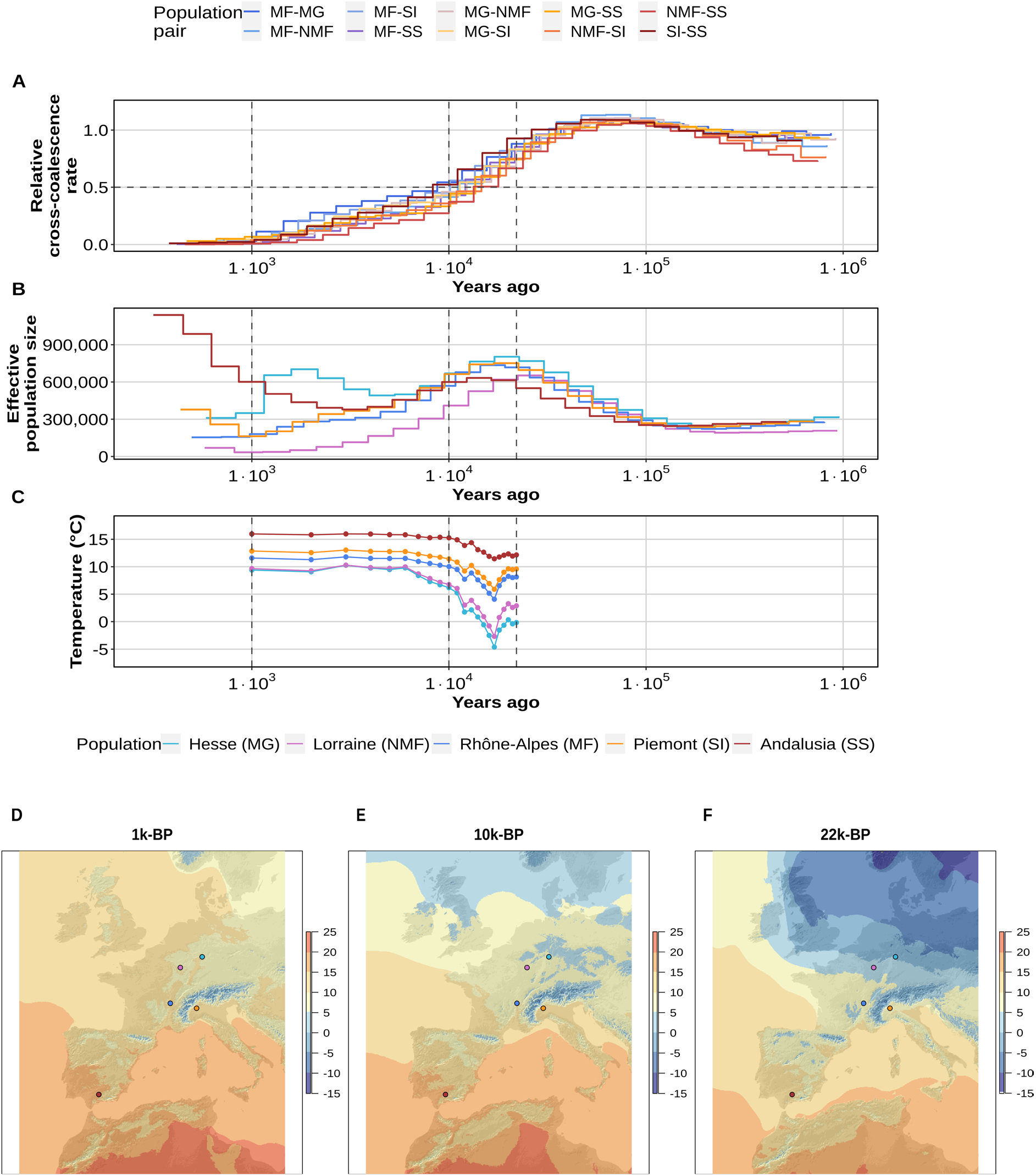
Comparison of the demographic history of *C. riparius* populations to available paleoclimate models. A) The relative cross-coalescence rate of the populations over the years reaching into the past estimated by the MSMC2 model. The population pairs are indicated by colour, as shown in the legend on top. B) Inferred effective population size over years reaching into the past estimated by MSMC2. Colour indicated by legend at the bottom. C) Annual mean temperature every thousand years for the separate sampling locations. D-F) Maps of Europe showing the spatial temperature pattern across Europe 1k-BP, 10k-BP and 22k-BP. Temperature (°C) is indicated by the gradient bar on the right. Dots refer to the sampling locations of the populations with corresponding colour code.

Analysing the estimated population demography to the millennial temperature intervals revealed that the zenith of population sizes occurred prior to a temperature decline about 16,000 years ago. The lowest temperature, with an annual mean temperature of-4.6 °C, was registered 17,000 years ago at the site where the MG population is located today (Figure 2C). Subsequently, temperatures increased, interrupted only by a minor decrease in temperature 12,000 years ago. The temperature increment slowed down from 7,000 years till 1,000 years ago.

### The genomic influence of the Cla-element on the recombination landscape

To gain insights into the topology of the recombination landscape of the *C. riparius* populations, we estimated recombination rates (ρ) of the populations with four individuals each in 10 kb windows along the genome using the tool iSMC (Supplementary Table S5). For each chromosome, mean recombination rates were calculated based on the matching windows for each population and then compared to the population-specific position of *Cla*-elements (Figure 3). The variation in recombination rate across chromosomes was heterogeneous among populations, especially the populations from Lorraine (NMF), Rhône-Alpes (MF) and Piemont (SI) which revealed a lot of intrapopulation variation.

**Figure 3:**
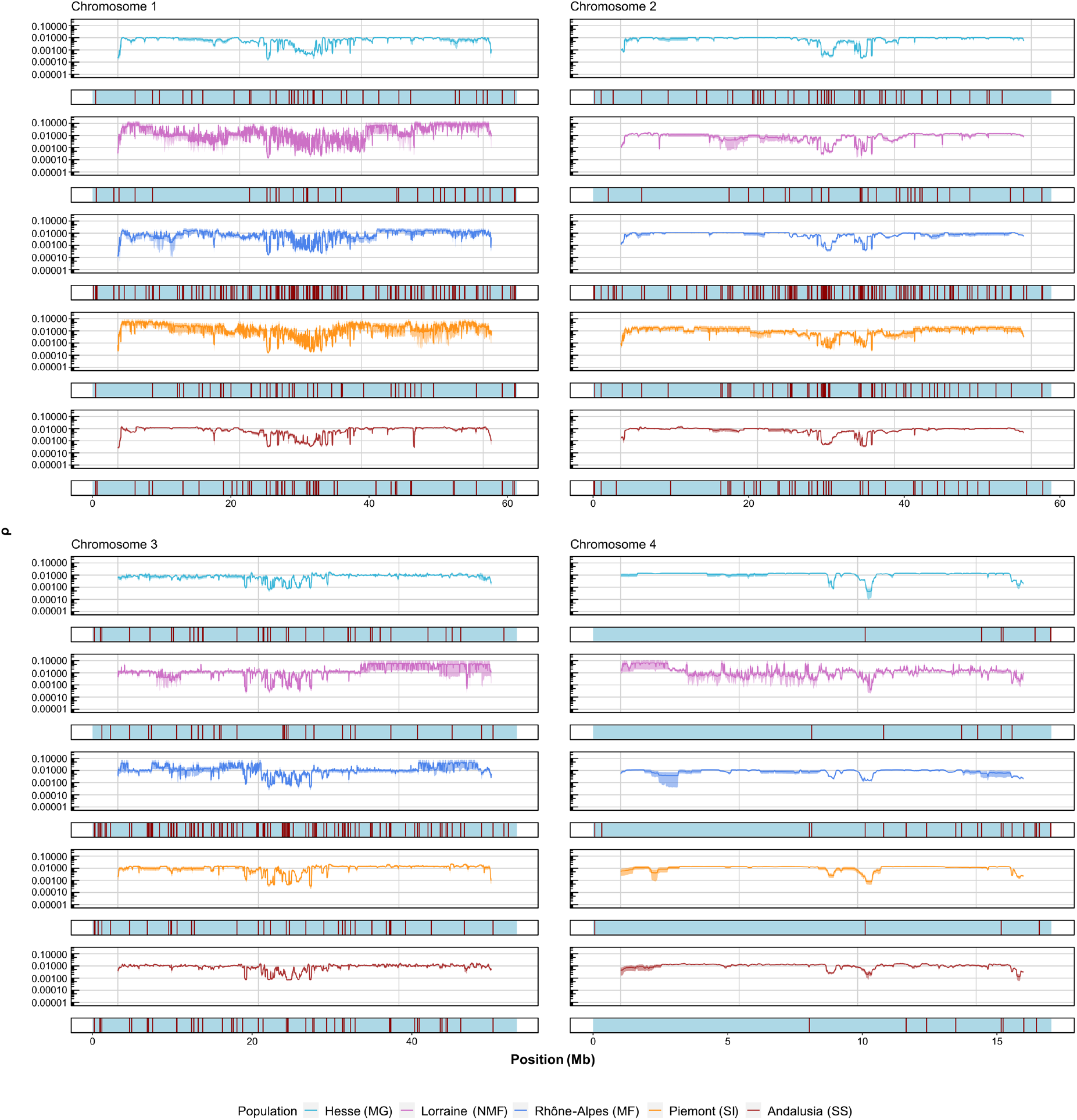
Mean recombination rate (ρ) (n=4) with confidence intervals in 10 kb windows on a log10-scale compared to the position of the *Cla*-elements along each chromosome, bottom of each chart. Position is given in Mb. Recombination rates and *Cla*-elements are shown separately for each population and colour-coded accordingly. Red lines mark the presence of a *Cla*-element.

Most *Cla*-elements were found on chromosome 1 and the fewest on the shortest chromosome 4. When summarising the total number of insertions (shared and unique) 441 insertions were identified. When looking at the populations independently, a total number of 188 insertions for the Andalusia (SS) population, 157 for the Piemont (SI) population, 148 for the Lorraine (NMF) population, 129 f or the Rhône-Alpes (MF) population and 107 for the Hesse (MG) population was detected.

To further investigate the relationship between the presence of *Cla*-elements and the local recombination landscape, we investigated the decay of ρ to the distance of the next Cla-element (Figure 4, Supplementary Figure S1, Supplementary Figure S2). For the cluster smaller than 500 bp there was a total of eight chromosomes of the analysed mean ρ with a moderate positive correlation (> 0.125) to the increasing distance of the *Cla*-element and for one we could find a strong positive correlation (> 0.25).

**Figure 4:**
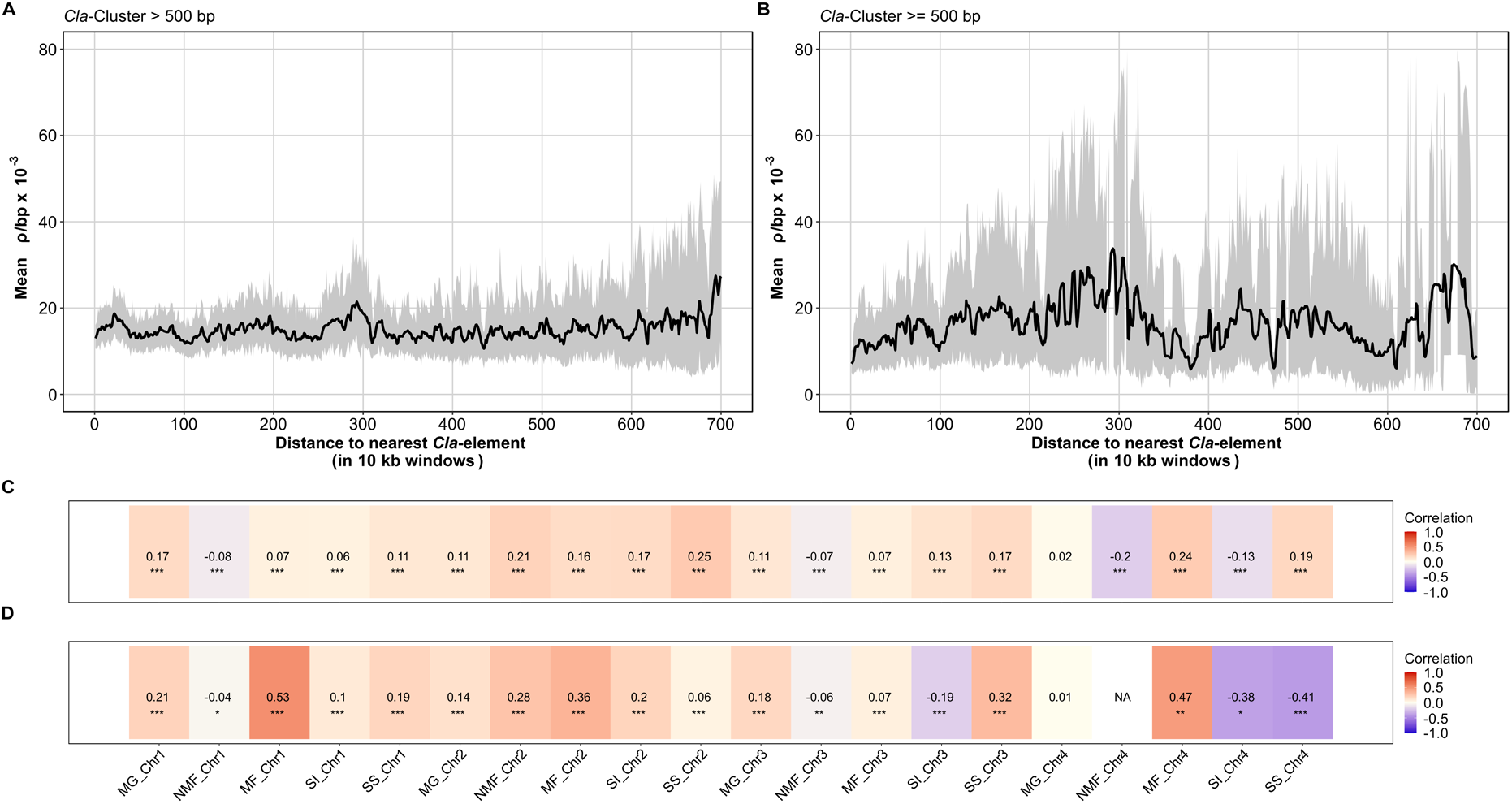
A-B) Mean recombination rate ρ/bp against distance to nearest Cla-element on chromosome 1. Mean values were calculated for each window based on each individual (n = 20). Displayed in grey are 100 bootstrap values calculated by resampling the mean to visualise its distribution. A) Cla-element cluster size <500 bp. B) Cla-element cluster size >=500 bp. C-D) Pearson correlation between recombination rate ρ and distance to next Cla-element. An asterisk indicates significance (* p < 0.05, ** p < 0.01, *** p < 0.001). Positive correlation indicates a higher recombination rate with a more distant Cla-element. C) Cla-element cluster size <500 bp. D) Cla-element cluster size >=500 bp. For these figures only complete windows of 10 kb were regarded. Cla-elements more distant than 7×10^6^ bp were excluded.

For clusters larger than or equal to 500 bp there was a moderate positive correlation and a strong positive correlation for five chromosomes each. However, not all chromosomes were positively correlated to higher recombination rates in regions with more distant *Cla*-elements (Figure 4A-B). For all clusters a highly variable mean ρ/bp x 10^-3^ could be identified compared to the distance to the nearest *Cla*-element supported by bootstrapping. As such, there is no consistent global pattern between recombination and the *Cla*-element, neither on the genome-wide nor on the level of particular chromosomes.

## Discussion

This study presents a novel genome assembly of chromosome-resolution for the emerging model organism *C. riparius*, a non-biting midge species also called harlequin fly. We test the impact of genome quality on the increase in resolution of demographic inferences by comparing estimates to a previous study. To reveal the genomic patterns that underly estimates of population history, we uncover the genome-wide landscape of recombination rates also looking at the potential role of a particular TE in these patterns. Over and above these novel genomic insights into the genomic landscape of *C. riparius*, these genomic resources will be more generally valuable for comparative studies on insect genomics (Blackmon et al. 2017), experimental population genomics (Foucault et al. 2019) and chromosome evolution (Shaikhutdinov & Gusev 2022).

### More ancient population history estimates could be inferred for C. riparius

In a previous study, Waldvogel et al. (2018) used a fragmented genome assembly for an initial inference of population history in *C. riparius,* applying multiple sequential Markovian coalescence (MSMC2). Many short scaffolds and the lack of information about their placement hindered the inference of recombination sites and thus only 17.34 % (30 scaffolds ≥ 100 kb) of the assembly were suitable for the analysis. With the new genome assembly, we were able to input 99.28 % of the genome to the analysis-only excluding ten unplaced scaffolds. The new assembly has a BUSCO (diptera_odb10) completeness of 99.28 % (Table 1) demonstrating the advantage of long-read sequencing coupled with Hi-C scaffolding (Guiglielmoni et al. 2022). Our results indicate the importance of high-quality genomes as the time range of demographic events was shifted significantly in the new inference (Figure 2A-B).

Waldvogel et al. (2018) estimated an informative time horizon between 150,000 and 351,000 generations in the past, while in our study this range was 6,100 to 7,400,000 generations (Supplementary Figure S3). This increase in resolution on the demographic history of *C. riparius* in European populations can at least partially be explained by the high accuracy of PacBio long-reads leading to more accurate assemblies with better coverage and contiguity in low complexity and repetitive regions (Pollard et al. 2018). The chromosome-resolution of the assembly also allowed the improvement of MSMC2 preconditions, i.e. restricting the analysis to scaffolds with a minimum length of 500 kb (Schiffels & Wang 2020).

The inference of the recent past is known to reach better resolution when more sequences are investigated (Schiffels & Wang 2020). Under the assumption that young haplotype blocks should have larger sizes, the improved genome quality additionally contributes to the resolution of the more recent population history (Stumpf & McVean 2003).The new genome assembly provides a contiguous overview of repetitive regions and enables access to them, which, in the past has been proven to be difficult with short-read sequencing as repetitive elements may not be mapped uniquely and as a result sequencing reads would stay ambiguous or collapsed with high coverage but poor mapping quality (Pollard et al. 2018).

Furthermore, when inferring population history, the accuracy of (short) sequences was demonstrated to be important (Sellinger et al. 2021). The likelihood of unmapped reads is higher in assemblies based on short-read sequencing which means that if a fragmented genome is not of high resolution in problematic regions, estimates of population history will be biased (Sellinger et al. 2021). Technical errors, like spurious SNP calling or incorrect detection of TEs have been shown to decrease the accuracy of population history estimates (Sellinger et al. 2021). These errors are more likely, and more difficult to control for, in fragmented genomes with low resolution of low complexity regions.

In summary, the more complete and chromosome-level assembly used here likely resulted in a more accurate estimation of population recombination (ρ) and a more robust inference of population history spanning a wider period and reaching far deeper into the past. Our study also benefitted from the availability of a species-specific mutation rate estimate (Waldvogel & Pfenninger 2021) that additionally contributed to the increased resolution of population history in *C. riparius*.

### Biogeography supports population demography

Accurate population demography models allow us to interpret and correlate a population’s history with the biogeographic history of its habitat. The temperature developments of the CHELSA traCE21k time-series dataset, covering the LGM up to 1,000 years before present (Karger et al. 2021), were compared to the geographical coordinates of the sampling sites of the five *C. riparius* populations (Figure 2C-D). Correlation of these two very different data types, allowed us to investigate whether paleoclimate models could have the potential to support sequence-based demographic estimations.

For our MSMC2 analysis we adapted the time segment pattern, approximated the time to the most recent common ancestor (tMRCA) of one population and trimmed outer values to account for overestimations of the most recent and most ancient time interval, resulting from false positive or negative signatures of recombination (Schiffels & Wang 2020). Our results suggest the origin of one ancestral population for the five investigated populations (Figure 2A) as has been proposed (Waldvogel et al. 2018). Whilst the split of the ancestral population has been proposed to have happened around 10,000 to 1,000 generations ago, our new estimates re-define this time frame. The admixture between the population shows a reduction between 500,000 to 10,000 generations ago and the rCCR reached a value of 0.5 at approximately 100,000 generations in the past. When multiplying these coalescence estimates with the population-specific generation time available for this multivoltine insect species (based on Oppold et al. 2016) the estimates were converted into years. This conversion defines the period of divergence of the populations between late Pleistocene and early Holocene (Stroeven et al. 2016) (Figure 2).

Starting from the Last Glacial Maximum (LGM,) between 22,000 and 17,000 years ago, the ice margins in Europe started to recede which led to an almost ice-free central Europe 16,000 years ago (Douda et al. 2014; Ehlers 1990; Stroeven et al. 2016). Two major climate events can be found in the temperature data (Figure 2C)-the Heinrich event (H1) around 16,800 years ago (Bond et al. 1992; Heinrich 1988; Hemming 2004) and the Younger Dryas around 12,000 years ago (Carlson 2010; Keigwin & Lehman 1994). Considering the biogeographic history of central Europe, it seems plausible that these climatic changes have contributed to the decreasing effective population size in the ancient population of *C. riparius*. The increase in temperature in the late Pleistocene might have led to a first dispersal of the midges as more habitats became available (Oppold et al. 2016). Both, the potential expansion across an increased habitat space, as well as the two cooling events are likely to have contributed to the decrease of effective population size of the ancient population. The decrease in effective population size and potential dispersal (see also in Waldvogel et al. 2018) might have also led to a reduced admixture, finally leading to a split of populations. As such, climate data could be used as an observational measure to support the demographic history estimation of *C. riparius* in Europe.

### A higher resolution of the recombination landscape and genome-wide patterns of repetitive elements is an advantage of high-quality assemblies

Advanced genome assemblies offer improved coverage of repetitive regions, unveiling a novel perspective on genome dynamics.

Based on these advances we were able to estimate population-specific recombination landscapes (Figure 3). For each chromosome, five different landscapes could be identified and for each population the mean recombination rate was based on the recombination landscape of four individuals. Certain regions of low recombination were shown to match between the populations – this could indicate lower recombination in the range of centromeres as generally expected (Choo 1998). The cause for the heterogenous pattern of recombination (especially the population Lorraine (NMF), Rhône-Alpes (MF) and Piemont (SI)) could be genomic markers, which led us to the initial study on the effect of the transposable *Cla*-element on the recombination landscape (Kent et al. 2017). As potential driver of population divergence, the *Cla*-element was of special interest (Oppold et al. 2017).

Genomic markers like microsatellites, TEs, telomeres, centromeres or heterochromatin-rich regions are known influencers of the dynamics within a genome (Kent et al. 2017; Sellinger et al. 2023; Oppold et al. 2017; Bascón-Cardozo et al. 2024), as they strongly influence the genomic landscape of an organism. They also should have a direct effect on the calculations done by models like the ones used to infer population history estimates. A recently published study (Sellinger et al. 2023) suggests an improvement in the reliability of population history estimates through incorporation of additional markers beyond SNP data, as demonstrated by more accurate estimates through the addition of Single Methylated Polymorphisms (SMPs) as markers. The authors propose that the addition of multiple genomic or epigenomic markers in population genomic models might aid future studies when sample number is limited, i.e. in most non-model species. Due to the complex dynamics of genomes, it is reasonable to assume that some genomic markers might be more fitted for one taxon than another. If looking for a genomic marker which could aid in explaining genome evolutionary processes in *C. riparius*, a promising candidate would be the transposable *Cla*-element. This element has been suggested to play a role in the speciation within the genus as it is only found in the centromeric regions of *C. piger* but spreads across chromosomal arms in *C. riparius* (Hankeln & Schmidt 1987). It has been shown that it is a driver for population divergence in *C. riparius* and only 25 % of the *Cla*-insertions have been found to be shared among all populations investigated, showing that there are many population-specific insertions (Oppold et al. 2017). These population specific insertions also become apparent in the analysis with MELT reported here, where the population of Andalusia (SS) is showing the highest number of *Cla*-insertions. As the proportion of shared *Cla*-elements is limited, an ongoing or recent activity in this TE is probable (Oppold et al. 2017). Overall, the incorporation of regions that were challenging to sequence in the past have the potential to enhance population genomic studies. Such regions may evolve at a different evolutionary speed. This underscores the necessity for highly resolved genome assemblies when studying the population dynamics of non-model organisms.

### Recombination rate ρ and its correlation to the next *Cla*-element

The *Cla*-element has an influence on chromosomal breaks in regions with heterochromatin (Bovero et al. 2002), as such, it seems likely that it has a general effect on recombination variation (Oppold et al. 2017). Ectopic recombination can serve as a selective mechanism against TEs but proves to be a fitness disadvantage if it is occurring in an organism (Kent et al. 2017). Ectopic recombination is generally associated with genomic instability compared to meiotic recombination as a highly regulated process (Lam & Keeney 2015; Pâques & Haber 1999; Hastings 2010). Recombination has a huge influence on the genetic diversity of a species and the length of haplotype blocks divided through recombination events gives insights on the past effective population size. Thus, investigating recombination is crucial to resolve Lewontin’s Paradox (Stumpf & McVean 2003; Charlesworth & Jensen 2022; Buffalo 2021). In addition, it has been demonstrated that TEs might modulate the local recombination landscape; some TEs seem to co-localize with recombination hotspots in humans (McVean 2010) and other TEs seem to accumulate in low recombining regions (Kent et al. 2017). Generally, TE density is suggested to correlate negatively with recombination, because selection against ectopic recombination and gene disruption can lead to accumulation of TEs in low recombining regions (Kent et al. 2017). Regulations of TEs might also influence local recombination rates (Kent et al. 2017). As such, it is difficult to disentangle cause and effect: are TEs actively shaping recombination variation or is it the reversed effect caused by the silencing of these elements (Kent et al. 2017)?

To investigate this in *C. riparius*, we compared the recombination landscape with the *Cla*-element pattern along the four different chromosomes (Figure 3 and 4). Most of the populations displayed a positive correlation, however there was no consistent pattern. A positive correlation would indicate a higher recombination rate if the next *Cla*-element is more distant. However, there were different levels of correlation between chromosomes of the different populations. Accordingly, there is no global pattern that connects the recombination landscape with the genome-wide distribution for *Cla*-elements in the genome of *C. riparius* even though the element has been described to drive population divergence (Oppold et al. 2017). However, it cannot be ruled out that the high number of *Cla*-insertions in the genome are convoluting the global analysis. The high number might be of varying age, and it is possible that older insertions already led to suppressed recombination and newer insertions might be too recent to find a co-evolving effect (Kent et al. 2017). If the transposition of the *Cla*-element is still active (Oppold et al. 2017) these signals might be covered. It could be also possible that it needs a certain number of insertions in close proximity, to detect reduced recombination to avoid ectopic recombination (Kent et al. 2017). The so-called insertion bias describes how TEs might be more commonly found in heterochromatin regions which indicates that TEs are expected to occur more frequently in low-recombing regions (Kent et al. 2017). However, there are also indicators for differential and purifying selection against TEs to select against ectopic recombination or gene disruption (Kent et al. 2017). There is not only a selective force due to direct fitness effects, but hosts appear to have evolved TE silencing mechanisms to decrease negative fitness effects (Kent et al. 2017). Natural selection and genetic drift could drive TEs in low recombining regions, but TE silencing could actively suppress recombination to repress transposition and ectopic recombination to avoid harmful fitness effects (Kent et al. 2017). As such, TE silencing could be an important modulator of local and large-scale recombination rates and might also explain differences in recombination landscape of con-specific populations (Kent et al. 2017; Mérel et al. 2020). Here, we could not confirm a higher occurrence of the *Cla*-element in low recombining regions. This might be explained by the theory of an ongoing transposition suspected for *Cla*-elements (Oppold et al. 2017). Additionally, there could be little to no regulation of this TE based on characteristics (Oppold et al. 2017). If there are regulatory processes controlling transposition of TEs in *C. riparius*, they might influence local recombination rates. It can be noted that a differential pattern of recombination can be observed in those chromosomes with many *Cla*-elements (Figure 3). It might be useful to assess the age of TEs and proportion of TEs that are silenced and compare them to the local recombination rates as suggested by Kent et al. (2017)-with this a better understanding of local recombination variation could be obtained. Possible further investigations through comparative genomics might offer insights in resolving this question. Investigating this mechanism could create new perspectives in the genome evolution of *C. riparius*, especially considering the involvement of the *Cla*-element in the speciation process.

### Conclusions

Based on a novel genome assembly at a chromosome scale, population history estimates of improved accuracy were obtained. We could show a shift in the coalescence estimates using MSMC2 compared to the previous study and could match these new results to past climate events. As recombination variation can highly influence population history estimates, the recombination landscape of *C. riparius* should be investigated. Population-specific recombination landscapes were successfully estimated giving a first insight into recombination variation of the five different European populations. With a genome assembly of increased resolution more regions of the genome including repetitive elements, like the *Cla*-element, could be investigated. The potential impact of *Cla* insertions on the recombination landscape was explored but no global pattern could be seen. We could show that the recombination landscape of *C. riparius* seems to be more stable regarding the impact of repetitive elements on genome integrity than previously assumed.

## Materials and Methods

### Data

The reference genome of *Chironomus riparius* provided by Rothamsted Research (West Common, Harpenden, United Kingdom) was assembled from a single female individual (German origin, procured Innovative Environmental services (IES) Ltd, Switzerland). DNA was extracted using MagAttract and 450ng was sequenced with PacBio HiFi using one SMRT cell and Hi-C sequencing using Illumina. The genome contains four chromosomes and ten scaffolds with a size of 192 Mb (NCBI accession number: PRJEB47883). Five different *C. riparius* populations, which were directly taken from their natural habitat, were studied. These are the same populations as were investigated in a previous study (Waldvogel et al. 2018). The origin of the different populations was from Rhône-Alpes (MF) and Lorraine (NMF) in France, Hesse in Germany (MG), Piedmont in Italy (SI) and Andalusia in Spain (SS). Twenty individual resequencing datasets were downloaded from the European Nucleotide Archive (ENA: 150-bp paired-end, Illumina sequencing data; Project number PRJEB24868). For each population four individuals were available. Of the datasets, the forward and reverse reads were already trimmed.

### Genome assembly and annotation

Hifiasm (Cheng et al. 2022, 2021) was used to assemble the PacBio HiFi data, then with Juicer (Durand et al. 2016) and 3d-dna (Dudchenko et al. 2017) the Hi-C data was used for chromosome level assembly. Haplotigs were removed using purge_haplotigs (Roach et al. 2018). Unmapped reads were mapped back to the original assembly to check for missing sequences and incorporated into the final assembly. Manual curation was done to bring the genome together and check for miss-assemblies. To assess the general quality of the genome assembly the software Blobtoolskits (v2.6.5, Challis et al. 2020) was utilized and an analysis for BUSCO completeness was performed (v5.3.2, Manni, Berkeley, Seppey & Zdobnov 2021; Manni, Berkeley, Seppey, Simão, et al. 2021; Simão et al. 2015).

A public RNA-seq transcriptome was assembled (BUSCO Insecta: C:94.7%[S:53.7%,D:41.0%],F:0.4%,M:4.9%) and used in the Maker2 (Holt & Yandell 2011) annotation pipeline with trained Augustus (Stanke et al. 2008) and Genemark (Borodovsky & McIninch 1993) gene predictors. Data used included: PRJEB15223 (Larvae), PRJNA166085 (egg ropes, all four larval stages, pupae and male and female adults, larvae exposed to different concentrations of several model toxicants) PRJNA229141 (anterior and posterior early embryo), PRJNA675286 (larvae-transition metal oxide exposure). PASA (Haas et al. 2003) was used to update the gene models to add UTR, correct existing models and add isoforms. Non-coding RNA was annotated using Infernal (v1.1.4, Nawrocki & Eddy 2013).

A Pfam genomic track was created by converting to six reading frames and applying hmmer (Finn et al. 2015) to identify loci of interest. Using this information, UDP, P450, ABC and IRAC gene models were found and curated using mapped RNA-seq and a Maker gene annotation.

Two Endosymbionts were assembled which included an unknown *Enterobacter* sp. (1,661,850 bp) and *Wolbachia* sp. (559,667 bp).

### Data pre-processing

Read quality was checked with FastQC (v0.11.9, Andrews 2010) and MultiQC (v1.12, Ewels et al. 2016). All pre-processing steps were performed according to Waldvogel et al. (2018). The trimmed reads were mapped separately to our reference genome using the tool bwa mem (-M-R’@RG\tID:$Population\tSM:$Individual\tPL:ILLUMINA’, v0.7.17, Li 2013). Low quality alignments were removed using samtools (-q 30-f 0×0002-F 0×0004-F0×0008, v1.13, Li et al. 2009) and to remove duplicates PicardTools (VALIDATIONSTRINGENCY SILENT-REMOVEDUPLICATES true, v2.26.10, Broad Institute 2018) was utilised. Mapping statistics were obtained using Qualimap (v2.2.2d, Okonechnikov et al. 2016). Further details on all analysis steps can be found on GitHub: https://github.com/lpettrich/Crip_Recombination_PopHistory_Cla_2024.

### Variant calling

Variant calling and phasing were performed as suggested in the MSMC2 workflow (Schiffels 2016) for the four chromosomes (99.28 % of the assembly). The reference genome was split by chromosome and mappability masks were created using SNPable (Li 2009), indicating mappable regions of the genome assembly. Following the standard workflow, variant calling was performed on the filtered bam-files of the samples using bcftools (v1.13, Li 2011) and the script bamCaller.py supplied by msmc-tools (Schiffels 2021) disregarding indels. This workflow generated VCF-and mask-files for each individual and chromosome which were necessary for generating the input files for MSMC2. Phasing was performed per chromosome utilising SHAPEIT4 (v4.2, Delaneau et al. 2019). Since no reference panel for *C. riparius* was available, we merged all VCF-files (bcftools merge) for phasing and separated them once again. To account for any missing information that is still contained in the unphased data, the phased and unphased VCF-files were merged while keeping the unphased data and replacing it with phased data where it was available. All multiallelic SNPs were discarded and only biallelic sites were kept. Using the obtained masking files and variant calls, multihetsep-files were generated using the script generatemultihetsep.py of the msmc-tools.

### Population demography

The generated multihetsep-files were used for MSMC2 (v2.1.3, Schiffels & Wang (2020)). Two populations were paired, resulting in a total of sixteen haplotypes (four diploid individuals per population) per dataset and a total of ten different population pairs that were later analysed in a cross-population analysis. The procedure of the cross-population analysis was to allow the first two MSMC2-runs that estimated the coalescence rate function within the population. Afterwards, an analysis across the populations was performed, selecting the population pairs. For the analysis the used time segment pattern was 1*3+1*2+22*1+1*2+1*3 and ambiguous sites were skipped. In the end the results were combined using the combineCrossCoal.py script from the msmc-tools. Overall, the obtained output per population included time and population size estimates, as well as the relative cross-coalescence rate (rCCR) which is a measure indicating the divergence of populations. Time and population size estimates were averaged per population and then scaled to real time and effective population size. Time estimates were converted into generations by dividing it through the mutation rate of 4.27 x 10^-9^ (Waldvogel & Pfenninger 2021). By multiplying it with the generation time (Waldvogel et al. 2018; Oppold et al. 2016) these converted coalescence times were converted into years. The effective population size was obtained by inverting the coalescence rate and dividing it by two-times the mutation rate. The first five entries, as well as the last entry were excluded to account for uncertainties in the analysis. Additionally, the mean haplotype length (MHL) was determined through the mean genome-wide heterozygosity and with regards to the switch error rate (SER). The SER of 2 % in *Drosophila melanogaster* (Bukowicki et al. 2016) was used, the same as in the previous study of Waldvogel et al. (2018). The mean heterozygosity was determined from the ratio of diallelic SNPs per number of records. These values were needed to approximate the time to the most recent common ancestor (tMRCA) of one population with the following formula: tMRCA= 1/(2∗r∗MHL). The population recombination rate (ρ) (based on Schmidt et al. 2020) was approximated to the recombination rate in units of meiosis per generation (r) using this formula from Peñalba and Wolf (2020): r=ρ/(2∗c∗Ne). In this context, c and Ne represented the organism’s diploidy and effective population size respectively. The effective population size (Oppold & Pfenninger 2017) was used for the calculation. To get an approximation of r in cM/Mb the gene map length of *D. melanogaster* of 287.3 cM (Comeron et al. 2012) was used as it should be comparable to *C. riparius* where the gene map length is not yet available. The mean value of r of 1.36 cM/Mb was then used to calculate tMRCA. This value was compared to the one using the recombination rate of *D. melanogaster* of 2.1 cM/Mb (Mackay et al. 2012) which is the same value as used in Waldvogel et al. (2018).

### Paleoclimate data

Results of the MSMC2 model of *C. riparius* were further compared to paleoclimate temperature data. Therefore, twenty-two bio1-datasets of the CHELSA-TraCE21k climate time-series were downloaded from CHELSA (Karger et al. 2021). Thus, the CHELSA-TraCE21k climate data provides information for the last 22,000 years before present (years BP) which referred to the Last Glacial Maximum (LGM). As such, contained the bio1-datsets annual mean temperatures and the here used twenty-two datasets (Supplementary Table S2) included timepoints from 1,000 years BP up to 22,000 years BP and were retrieved in steps of one thousand years (i.e., millennial time series).

### Recombination landscape

To reconstruct the recombination landscape of *C. riparius* we used the tool iSMC (Barroso et al. 2019) and the provided scripts of Robinson et al. (2021). This tool uses SMC models to infer population history and spatial variation in recombination rates. The artificial phased VCF-files were used as input files. The chromosomes were analysed separately per individual. For each analysis an iSMC option file needed to be provided where we used the standard recommendations from the iSMC documentation (optimize = true, decode = true, decodebreakpointsparallel = false, num-berrhocategories = 5, numberintervals = 40, functiontolerance = 1e-4, fragmentsize = 3000000). To obtain recombination rates (ρ) and information on tMRCA we used the iSMC mapper with a bin size of 10 kb and 1 Mb.

### Transposable elements

Sensitive soft masking of repeats on the genome was performed with RepeatMasker (v4.1.1, Smit et al. 2015) using a custom TE-library by Vladimir Kapitanov which was modified by adding TE-entries of Oppold et al. (2017). A cutoff score of masking repeats of 250 bp and the engine rmblast (-s-xsmall-cutoff 250-u-gff-pa 10-lib $LIB-dir $DIR-enginermblast $GENOME) was chosen for the RepeatMasker analysis. The output file was converted to BED format using RM2Bed.py script supplied by RepeatMasker. Additionally, an annotation of the Cla-element population-wise was conducted using MELT (v2.2.2, Gardner et al. 2017). The necessary mei.zip file for the Cla-element was created following the instructions of the MELT documentation. To find the insertions we added the MQ tag using samblaster to our sam-files and used the unfiltered but sorted bam-files as input for MELT-SPLIT. Data for each of the four different analyses runs were obtained for each population and for all populations together. To visualise the insertions of the *Cla*-element in the genome of each population compared to their inferred recombination rates, R (v4.2.1, R Core Team 2020) and R-Studio (v2022.02.0+433, RStudio Team 2020) was used together with several R packages, like tidyverse (v2.0.0, Wickham et al. 2023), patchwork (v1.1.3, Pedersen 2024) or cowplot (v1.1.1, Wilke 2020). A detailed list of the R packages can be found in the Supplementary Table S1. To test whether the occurrence of *Cla* insertions is correlated to recombination events, we used the closest function of bedtools (v2.31.0, Quinlan & Hall 2010), according to Robinson et al. (2021). R-Studio was used to create plots showing the recombination rate in relation to the nearest Cla-element. These decay plots were created using the distance of the next Cla-element to a recombination event. *Cla*-clusters with sizes smaller than 500 bp were investigated separately from *Cla*-Clusters larger or equal to 500 bp. The obtained measures were tested with the Pearson correlation coefficient using the R package stats and results were then visualised in a heatmap. Pearson correlation coefficients were calculated for each chromosome of each population (mean values were calculated from the four individuals per population) to investigate whether the distance of the *Cla*-element is correlated to the amplitude of ρ. Smaller and larger cluster were again analysed independent from each other. For the creation of loops in R and similar simplifications of the scripts ChatGPT was utilised (OpenAI 2024).

## Acknowledgments

We thank Dr. Vladimir Kapitanov from the LOEWE Center for Translational Biodiversity Genomics, Frankfurt am Main, Germany for support with the development and nomenclature of the TE library that was used for the TE annotation of the genome. We thank Dr. Philipp Schiffer and Laura I. Villegas for their thoughtful comments on this manuscript. AMW acknowledges the funding of her junior professorship in the “Bund-Länder Programm” of the German Federal Ministry of Education and Research (BMBF). This work has additionally been funded as part of subproject B08 in the Collaborative Research Cluster 1211 (DFG grant number 268236062). We furthermore acknowledge the Regional Computing Center of the University of Cologne (RRZK) for providing computing time on the DFG-funded [Funding number INST 216/512/1FUGG] High Performance Computing (HPC) system CHEOPS as well as support.

## Author Contributions

L.C.P. and A.M.W. perceived the study. L.C.P. performed the data analysis and drafted the manuscript. A.M.W. contributed to writing and supervised the work. L.M.F. and R.K. generated and provided the genome assembly. All authors approved the final version of the manuscript.

## Data Availability

The genome assembly can be downloaded at ENA (accession PRJEB47883). The Illumina sequences of the five populations have been published under Waldvogel et al. (2018) and trimmed reads can be accessed at ENA (accession PRJEB24868). Scripts are available at the GitHub repository https://github.com/lpettrich/Crip_Recombination_PopHistory_Cla_2024.

## Notes

### Competing Interest Statement

The authors have declared no competing interest.

