## Supplement for "The interplay of recombination landscape, a transposable element and population history in European populations of *Chironomus riparius*"

Author details:

1 Institute of Zoology, University of Cologne, Cologne, Germany

2 Rothamsted Research, Harpenden, Hertfordshire, United Kingdom

*Authors for Correspondence:

Laura Chiara Pettrich, Institute of Zoology, University of Cologne, Cologne, Germany,

and

Ann-Marie Waldvogel, Institute of Zoology, University of Cologne, Cologne, Germany,

### Supplementary Materials

Supplementary Table S1: List of tools and R packages used with citations.

| | Tools | Version | Citation |
|---|---|---|---|
| **Software** | samtools | 1.13 | Li et al. (2009) |
| | bcftools | 1.13 | Li et al. (2009) |
| | bwa | 0.7.17 | Li (2013) |
| | Trimmomatic | 0.39 | Bolger et al. (2014) |
| | Picard Tools | 2.26.10 | Broad Institute (2018) |
| | FastQC | 0.11.9 | Andrews (2010) |
| | MultiQC | 1.12 | Ewels et al. (2016) |
| | Qualimap | 2.2.2d | Okonechnikov et al. (2016) |
| | bedtools | 2.31.0 | Quinlan & Hall (2010) |
| | shapeit4 | 4.2 | Delaneau et al. (2019) |
| | MSMC2 | 2.1.3 | Schiffels & Wang (2020) |
| | msmc-tools | - | Schiffels (2021) |
| | SNPable | - | Li (2009) |
| | iSMC | - | Barroso et al. (2019) |
| | RepeatMasker | 4.1.1 | Smit et al. (2015) |
| | MELT | 2.2.2 | Gardner et al. (2017) |
| | blastx | 2.12.0 | Camacho et al. (2009) |
| | BUSCO | 5.3.2 | Manni et al. (2021), Simão et al. (2015) |
| | BlobToolsKit | 2.6.5 | Challis et al. (2020) |
| | Rstudio | 2022.02.0+433 | RStudio Team (2020) |
| | R | 4.2.1 | R Core Team (2020) |
| **R packages** | ggplot2 | 3.3.6 | Wickham (2016) |
| | dplyr | 1.1.4 | Wickham et al. (2023) |
| | tidyverse | 2.0.0 | Wickham et al. (2019) |
| | raster | 3.5-15 | Hijmans & van Etten (2012) |
| | egg | 0.4.5 | Auguie (2019) |
| | grid | 4.2.1 | R Core Team (2020) |
| | patchwork | 1.1.3 | Pedersen (2024) |
| | cowplot | 1.1.1 | Wilke (2020) |
| | scales | 1.2.1 | Wickham & Seidel (2022) |
| | gridExtra | 2.3 | Auguie (2017) |
| | reshape2 | 1.4.4 | Wickham (2007) |
| | RColorBrewer | 1.1-3 | Neuwirth (2022) |
| | sf | 1.0-14 | Pebesma (2018), Pebesma & Bivand (2023) |
| | rnaturalearth | 0.3.4 | Massicotte & South (2023) |
| | naturalearthdata | 0.1.0 | South (2017) |
| | rgeos | 0.6-4 | Bivand & Rundel (2023) |
| | maptools | 1.1-8 | Bivand & Lewin-Koh (2023) |
| | rgdal | 1.6-7 | Bivand et al. (2023) |
| | ggpubr | 0.6.0 | Kassambara (2022) |
| | gridGraphics | 0.5-1 | Murrell & Wen (2020) |

Supplementary Table S2: Time periods from the CHELSA-TraCE21k dataset (Karger et al. 2021) used in this study. Further information can be found on the CHELSA webpage.

| **Time ID** | **Start year** | **End year** | **k-BP** |
|---|---|---|---|
| 10 | 900 | 999 | 1 |
| 0 | -100 | -1 | 2 |
| -10 | -1100 | -1001 | 3 |
| -20 | -2100 | -2001 | 4 |
| -30 | -3100 | -3001 | 5 |
| -40 | -4100 | -4001 | 6 |
| -50 | -5100 | -5001 | 7 |
| -60 | -6100 | -6001 | 8 |
| -70 | -7100 | -7001 | 9 |
| -80 | -8100 | -8001 | 10 |
| -90 | -9100 | -9001 | 11 |
| -100 | -10100 | -10001 | 12 |
| -110 | -11100 | -11001 | 13 |
| -120 | -12100 | -12001 | 14 |
| -130 | -13100 | -13001 | 15 |
| -140 | -14100 | -14001 | 16 |
| -150 | -15100 | -15001 | 17 |
| -160 | -16100 | -16001 | 18 |
| -180 | -18100 | -18001 | 20 |
| -190 | -19100 | -19001 | 21 |
| -200 | -20100 | -20001 | 22 |

### Supplementary Results

Supplementary Table S3: Mapping statistics of resequencing data.

| **Sample** | **Number of reads** | **Mapped reads** | **Properly paired reads** | **Genome-wide mean coverage** | **Insert size median (bp)** | **GC-content (%)** |
|---|---|---|---|---|---|---|
| MF1 | 37512849 | 36680245 (97.78 %) | 33345658 (90.44 %) | 15.5912 | 297 | 32.61 |
| MF2 | 30925208 | 30305675 (98.00 %) | 27573498 (90.74 %) | 13.2659 | 298 | 32.15 |
| MF3 | 32731900 | 31546904 (96.38 %) | 28768950 (89.43 %) | 13.642 | 296 | 32.92 |
| MF4 | 32888448 | 32070040 (97.51 %) | 29402506 (90.99 %) | 16.6019 | 273 | 32.72 |
| MG2 | 29574915 | 29098966 (98.39 %) | 26464018 (91.07 %) | 12.4309 | 288 | 32.61 |
| MG3 | 29282686 | 28769324 (98.25 % ) | 26096752 (90.65 %) | 12.3216 | 298 | 32.63 |
| MG4 | 27560169 | 27107536 (98.36 %) | 24692740 (91.14 %) | 11.6459 | 289 | 32.55 |
| MG5 | 34723726 | 34098931 (98.20 %) | 31279922 (91.73 %) | 17.3902 | 272 | 33.45 |
| NMF1 | 39149237 | 38418718 (98.13 %) | 34846302 (90.58 %) | 19.2096 | 291 | 32.38 |
| NMF2 | 32848323 | 32332865 (98.43 %) | 29604680 (91.74 %) | 16.5641 | 279 | 33.07 |
| NMF3 | 29922750 | 29270955 (97.82 %) | 26781690 (91.13 %) | 15.2467 | 272 | 32.73 |
| NMF4 | 26926669 | 26460488 (98.27 %) | 24110414 (91.17 %) | 13.7049 | 277 | 32.71 |
| SI1 | 38628115 | 37887945 (98.08 %) | 34543616 (91.03 %) | 16.6717 | 294 | 32.58 |
| SI2 | 43087974 | 41349691 (95.97 %) | 37532316 (88.67 %) | 17.0428 | 288 | 32.68 |
| SI3 | 33478250 | 31731906 (94.78 %) | 28829146 (87.57 %) | 14.0101 | 300 | 32.67 |
| SI4 | 31364052 | 26697695 (85.12 %) | 24160270 (78.26 %) | 13.4785 | 287 | 32.23 |
| SS1 | 31635833 | 31148191 (98.46 %) | 28308742 (91.09 %) | 13.4556 | 298 | 32.62 |
| SS2 | 40525533 | 39859702 (98.36 %) | 36349244 (91.36 %) | 16.8263 | 289 | 31.91 |
| SS3 | 36668799 | 36066090 (98.36 %) | 32808088 (91.09 %) | 15.7296 | 288 | 31.94 |
| SS4 | 29302810 | 28821648 (98.36 %) | 26208916 (91.08 %) | 14.7396 | 285 | 33.12 |

Supplementary Table S4: Final diallelic SNP count per chromosome and individual of *Chironomus riparius*. Entries in mulithetsep files were used as input in the models of MSMC2 and iSMC.

| Chromosme | Sample | SNP count | Entries in multihetsep |
|---|---|---|---|
| 1 | MF1 | 902,195 | 183,734 |
| 1 | MF2 | 902,289 | 198,319 |
| 1 | MF3 | 901,883 | 129,659 |
| 1 | MF4 | 902,269 | 200,968 |
| 1 | MG2 | 902,062 | 218,160 |
| 1 | MG3 | 901,970 | 194,808 |
| 1 | MG4 | 902,245 | 232,985 |
| 1 | MG5 | 901,889 | 165,313 |
| 1 | NMF1 | 901,953 | 101,194 |
| 1 | NMF2 | 901,811 | 108,234 |
| 1 | NMF3 | 902,160 | 171,242 |
| 1 | NMF4 | 902,100 | 181,757 |
| 1 | SI1 | 902,294 | 204,944 |
| 1 | SI2 | 902,233 | 170,877 |
| 1 | SI3 | 902,079 | 195,934 |
| 1 | SI4 | 902,043 | 173,416 |
| 1 | SS1 | 902,274 | 225,600 |
| 1 | SS2 | 902,438 | 256,088 |
| 1 | SS3 | 902,255 | 223,891 |
| 1 | SS4 | 902,048 | 207,688 |
| 2 | MF1 | 913,108 | 228,105 |
| 2 | MF2 | 913,238 | 243,861 |
| 2 | MF3 | 912,982 | 221,618 |
| 2 | MF4 | 913,092 | 198,876 |
| 2 | MG2 | 913,102 | 245,326 |
| 2 | MG3 | 912,927 | 215,805 |
| 2 | MG4 | 912,954 | 239,476 |
| 2 | MG5 | 912,838 | 207,184 |
| 2 | NMF1 | 913,117 | 175,957 |
| 2 | NMF2 | 913,040 | 195,278 |
| 2 | NMF3 | 912,912 | 182,349 |
| 2 | NMF4 | 912,902 | 184,483 |
| 2 | SI1 | 913,081 | 219,691 |
| 2 | SI2 | 913,023 | 197,655 |
| 2 | SI3 | 912,842 | 157,342 |
| 2 | SI4 | 913,201 | 267,170 |
| 2 | SS1 | 912,946 | 213,785 |
| 2 | SS2 | 913,215 | 251,323 |
| 2 | SS3 | 913,209 | 258,693 |
| 2 | SS4 | 912,950 | 217,469 |
| 3 | MF1 | 786,653 | 102,172 |
| 3 | MF2 | 786,865 | 165,753 |
| 3 | MF3 | 786,770 | 176,347 |
| 3 | MF4 | 786,893 | 183,199 |
| 3 | MG2 | 786,842 | 192,167 |
| 3 | MG3 | 786,771 | 191,009 |
| 3 | MG4 | 786,804 | 148,405 |
| 3 | MG5 | 786,641 | 160,750 |

| Chromosme | Sample | SNP count | Entries in multihetsep |
|---|---|---|---|
| 3 | NMF1 | 786,687 | 91,689 |
| 3 | NMF2 | 786,703 | 127,623 |
| 3 | NMF3 | 786,785 | 156,371 |
| 3 | NMF4 | 786,815 | 172,260 |
| 3 | SI1 | 787,059 | 217,880 |
| 3 | SI2 | 787,032 | 210,968 |
| 3 | SI3 | 786,868 | 196,400 |
| 3 | SI4 | 786,907 | 187,919 |
| 3 | SS1 | 786,861 | 191,661 |
| 3 | SS2 | 787,109 | 222,640 |
| 3 | SS3 | 786,921 | 212,354 |
| 3 | SS4 | 786,701 | 180,613 |
| 4 | MF1 | 246,169 | 45,355 |
| 4 | MF2 | 246,126 | 45,305 |
| 4 | MF3 | 246,121 | 40,092 |
| 4 | MF4 | 246,198 | 56,983 |
| 4 | MG2 | 246,135 | 46,197 |
| 4 | MG3 | 246,171 | 50,311 |
| 4 | MG4 | 246,137 | 53,441 |
| 4 | MG5 | 246,187 | 52,156 |
| 4 | NMF1 | 246,032 | 12,405 |
| 4 | NMF2 | 246,125 | 34,388 |
| 4 | NMF3 | 246,198 | 51,683 |
| 4 | NMF4 | 246,130 | 35,438 |
| 4 | SI1 | 246,229 | 53,298 |
| 4 | SI2 | 246,200 | 50,040 |
| 4 | SI3 | 246,234 | 57,178 |
| 4 | SI4 | 246,218 | 59,853 |
| 4 | SS1 | 246,184 | 52,714 |
| 4 | SS2 | 246,201 | 58,280 |
| 4 | SS3 | 246,175 | 50,133 |
| 4 | SS4 | 246,138 | 46,671 |

Supplementary Table S5: Global mean of ρ for each chromosome of each population.

| Population | Chromosome | N | Mean ρ | SE low | SE high |
|---|---|---|---|---|---|
| MF | Chr1 | 24536 | 0.00915354 | 0.00908653 | 0.00922055 |
| MF | Chr2 | 23548 | 0.00845388 | 0.00842641 | 0.00848134 |
| MF | Chr3 | 21256 | 0.01575148 | 0.01557013 | 0.01593283 |
| MF | Chr4 | 6804 | 0.00833753 | 0.0082824 | 0.00839267 |
| MG | Chr1 | 24536 | 0.00694967 | 0.00692573 | 0.0069736 |
| MG | Chr2 | 23548 | 0.00827227 | 0.0082477 | 0.00829683 |
| MG | Chr3 | 21256 | 0.00805335 | 0.00802291 | 0.00808378 |
| MG | Chr4 | 6804 | 0.01204706 | 0.01199279 | 0.01210134 |
| NMF | Chr1 | 24536 | 0.03253499 | 0.03215883 | 0.03291115 |
| NMF | Chr2 | 23548 | 0.00923108 | 0.00919161 | 0.00927055 |
| NMF | Chr3 | 21256 | 0.02270239 | 0.02240101 | 0.02300376 |
| NMF | Chr4 | 6804 | 0.01738729 | 0.01699122 | 0.01778336 |
| SI | Chr1 | 24536 | 0.02170701 | 0.02149756 | 0.02191645 |
| SI | Chr2 | 23548 | 0.01144847 | 0.01137648 | 0.01152046 |
| SI | Chr3 | 21256 | 0.01129308 | 0.0112511 | 0.01133506 |
| SI | Chr4 | 6804 | 0.01118637 | 0.01112667 | 0.01124606 |
| SS | Chr1 | 24536 | 0.00832284 | 0.0082943 | 0.00835138 |
| SS | Chr2 | 23548 | 0.00803629 | 0.0080133 | 0.00805928 |
| SS | Chr3 | 21256 | 0.00911988 | 0.00909169 | 0.00914807 |
| SS | Chr4 | 6804 | 0.01074792 | 0.01069637 | 0.01079948 |

Supplementary Figure S1: Decay of ρ to the distance of the next Cla-element all chromosomes for clusters >= 500 bp. Mean values were calculated for each window based on each individual (n = 20). Displayed in grey are 100 bootstrap values calculated by resampling the mean to visualise its distribution. For these figures only complete windows of 10 kb were regarded. *Cla*-elements more distant than 7x106 bp were excluded.

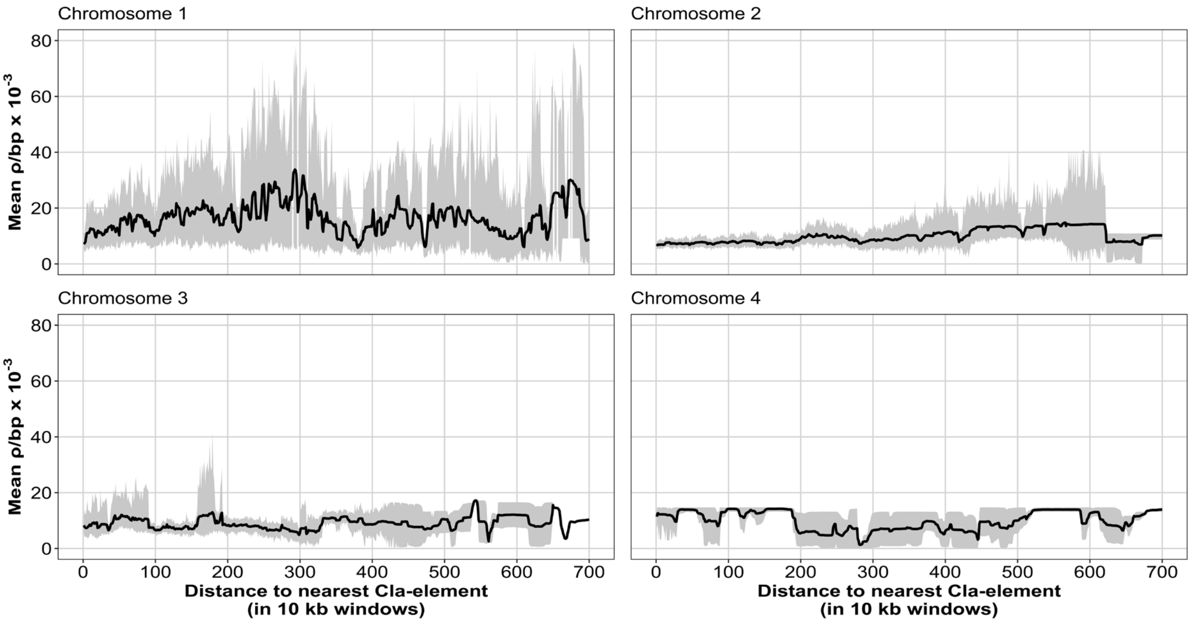

Supplementary Figure S2: Decay of ρ to the distance of the next Cla-element all chromosomes for clusters < 500 bp. Mean values were calculated for each window based on each individual (n = 20). Displayed in grey are 100 bootstrap values calculated by resampling the mean to visualise its distribution. For these figures only complete windows of 10 kb were regarded. Cla-elements more distant than 7x106 bp were excluded.

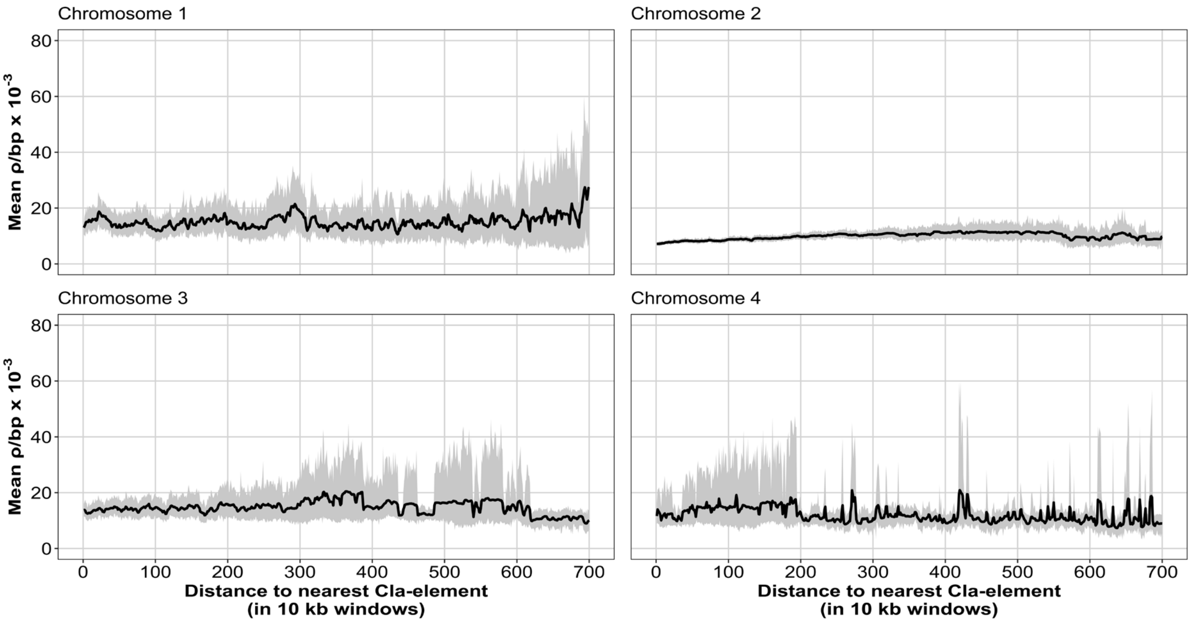

Supplementary Figure S3: MSMC2 analysis visualizing generations reaching into the past.

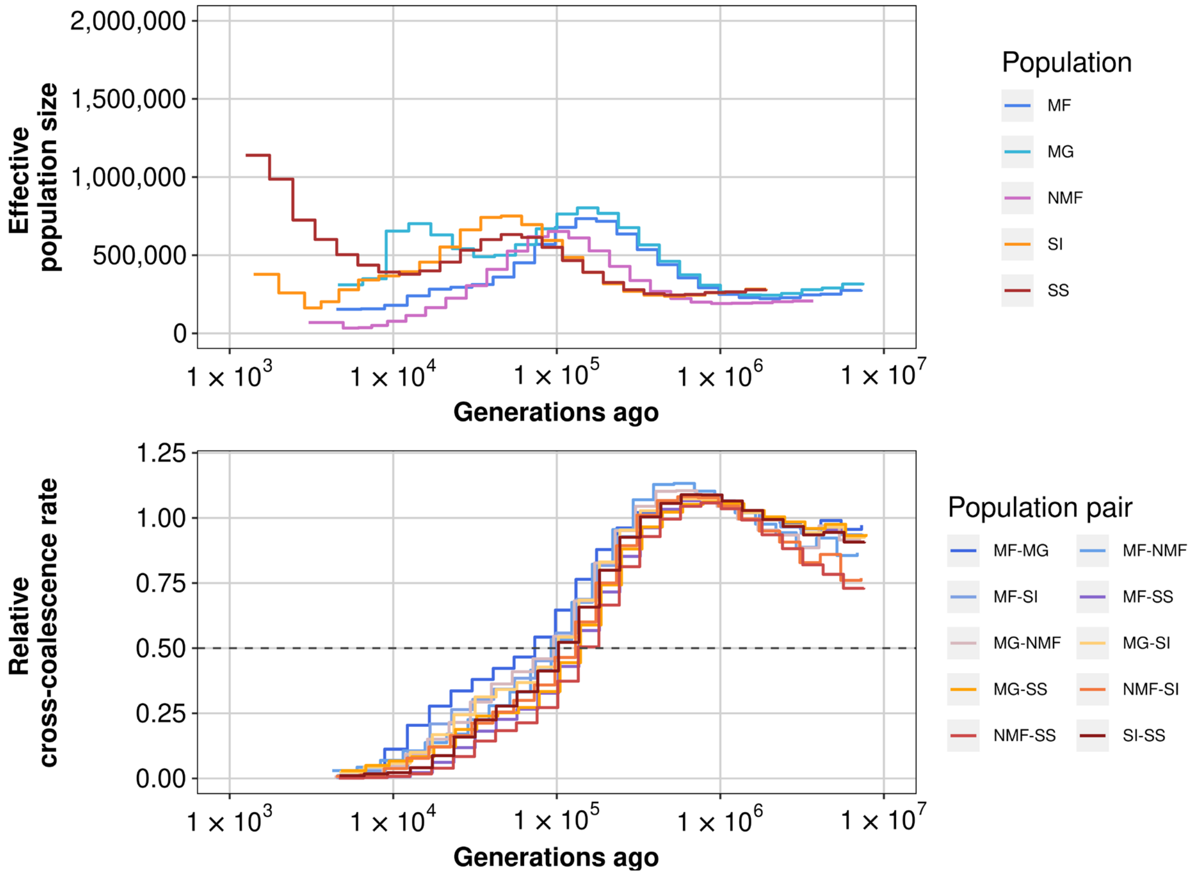


Supplementary Table S6: Summarising population specific parameters of *C. riparius* needed to convert recombination rates to estimate tMRCA in the MSMC2 analysis.

| **Population** | **Generations per year** (Oppold et al. 2016) | **Generation time** | **Recombination rate ρ (1/bp)** (Schmidt et al. 2020) | **Recombination rate r (cM/Mb)** | **Effective population size Ne** (Oppold & Pfenninger 2017) |
|---|---|---|---|---|---|
| MG | 7.85 | 0.1274 | 0.068 | 1.37 | 3570000 |
| NMF | 7.7 | 0.1299 | 0.040 | 0.73 | 3950000 |
| MF | 9.07 | 0.1103 | 0.058 | 1.21 | 3450000 |
| SI | 10.57 | 0.0946 | 0.060 | 1.31 | 3300000 |
| SS | 14.86 | 0.0673 | 0.066 | 2.20 | 2150000 |
| Mean | 10.01 | 0.0999 | | 1.36 | |
